# Competition for hosts modulates vast antigenic diversity to generate persistent strain structure in *Plasmodium falciparum*

**DOI:** 10.1101/406546

**Authors:** Shai Pilosof, Qixin He, Kathryn E. Tiedje, Shazia Ruybal-Pesántez, Karen P. Day, Mercedes Pascual

## Abstract

In their competition for hosts, parasites with antigens that are novel to host immunity will be at a competitive advantage. The resulting frequency-dependent selection can structure parasite populations into strains of limited genetic overlap. For *Plasmodium falciparum*–the causative agent of malaria–in endemic regions, the high recombination rates and associated vast diversity of its highly antigenic and multicopy *var* genes preclude such clear clustering; this undermines the definition of strains as specific, temporally-persisting gene variant combinations. We use temporal multilayer networks to analyze the genetic similarity of parasites in both simulated data and in an extensively and longitudinally sampled population in Ghana. When viewed over time, populations are structured into modules (i.e., groups) of parasite genomes whose *var* gene combinations are more similar within, than between, the modules, and whose persistence is much longer than that of the individual genomes that compose them. Comparison to neutral models that retain parasite population dynamics but lack competition reveals that the selection imposed by host immunity promotes the persistence of these modules. The modular structure is in turn associated with a slower acquisition of immunity by individual hosts. Modules thus represent dynamically generated niches in host immune space, which can be interpreted as strains. Negative frequency-dependent selection therefore shapes the organization of the *var* diversity into parasite genomes, leaving a persistence signature over ecological time scales. Multilayer networks extend the scope of phylodynamics analyses by allowing quantification of temporal genetic structure in organisms that generate variation via recombination or other non-bifurcating processes. A strain structure similar to the one described here should apply to other pathogens with large antigenic spaces that evolve via recombination. For malaria, the temporal modular structure should enable the formulation of tractable epidemiological models that account for parasite antigenic diversity and its influence on intervention outcomes.

**Significance:** Many pathogens, including the causative agent of malaria *Plasmodium falciparum*, use antigenic variation, obtained via recombination, as a strategy to evade the human immune system. The vast diversity and multiplicity of genes encoding antigenic variation in high transmission regions challenge the notion of the existence of distinct strains: temporally-persistent and specific combinations of genes relevant to epidemiology. We examine the role of human immune selection in generating such genetic population structure in the major blood-stage antigen of *Plasmodium falciparum*. We show, using simulated and empirical data, that immune selection generates and maintains ‘modules’ of genomes with higher genetic similarity within, than between, these groups. Selection further promotes the persistence of these modules for much longer times than those of their constituent genomes. Simulations show that the temporal modular structure reduces the speed at which hosts acquire immunity to the parasite. We argue that in *P. falciparum* modules can be viewed as dynamic strains occupying different niches in human immune space; they are thus relevant to formulating transmission models that encompass the antigenic diversity of the parasite. Our analyses may prove useful to understand the interplay between temporal genetic structure and epidemiology in other pathogens of human and wildlife importance.

## Introduction

The dynamic arms race between hosts and pathogens sets the stage for a selective advantage of rare variants that confer either immune protection to hosts or immune escape to pathogens, and a corresponding disadvantage of common ones. This frequency-dependent effect can act as a form of balancing selection and is a powerful force promoting high antigenic diversity and maintaining polymorphisms significantly longer than neutral drift. High diversity and temporal persistence of diversity at the gene level have been shown in numerous host-pathogen systems. Genes that encode pathogen resistance in hosts, such as the MHC [1], and those that underlie antigenic variation in parasites, such as the *var* multigene complex in the malaria parasite *Plasmodium falciparum*, display exceptional polymorphism compared to other functional genes. Several components of *var* genes are known to have originated millions of years ago and to be shared with closely-related species that infect apes [2].

In transmission systems, diversity and persistence are also relevant at a higher level of organization than that of individual genes. In particular, pathogen strains concern temporally-persistent combinations of genes related to infection, including those encoding antigens. The role of immune selection at this higher level of organization remains however poorly understood and documented, especially in pathogens whose antigenic variation involves vast diversity generated via recombination, within the genome or between different genomes, as is the case in several bacteria, protozoa and fungi [3]. Can strains exist and persist in such vast antigenic spaces? What is a strain in dynamic systems undergoing recombination at the level of both the genes themselves and the genomes they compose? A key characteristic signature of immune selection would involve the persistence of gene combinations over longer time scales than expected under neutrality [4], a hypothesis that remains to be examined despite its relevance for the existence and definition of strains themselves.

We address this temporal dimension by examining the role frequency dependence plays in maintaining gene combinations over time, in the highly diverse multicopy *var* gene family of *P. falciparum*. Frequency-dependent selection in pathogen systems is analogous to stabilizing competition in ecological communities [5, 6]. Competition can drive coexisting species to self-organize into clusters with limiting similarity along a niche or trait axis [7–10]. In epidemiology, strain theory had proposed that competition for hosts through cross-immunity, the cross-protection conferred by previous acquisition of immunity, can structure pathogen populations into temporally-stable sets of genetically distinct strains with limited overlap of antigenic repertoires [11]. Theoretical studies for low-to-medium genetic diversity predicted parasite strains with no, or limited, genetic overlap [12,13], including in the case of multicopy genes [14]. For *P. falciparum*, recent work allowing for realistic levels of genetic diversity comparable to that found in endemic high transmission regions of sub-Saharan Africa, resulted in a more complex similarity structure clearly distinguishable from patterns generated under neutrality but that can no longer be described by distinct clusters [15]. Deep sampling and sequencing of *var* gene isolates from asymptomatic human populations within a given time window or transmission season, have confirmed these non-random patterns that are also non-neutral, in the sense that they cannot be simply explained by stochastic extinction, immigration, and transmission in the absence of acquired immune memory and therefore, competition of parasites for hosts [15–17]. The lack of explicit consideration of the temporal dimension may be the reason why no apparent distinct clustering was identified, and is the motivation behind this work.

Analyzing the temporal dimension requires simultaneously tracking the population dynamics of genomes and their genetic relationship in a hyper-diverse system. An additional challenge is allowing for evolution via recombination. Here, we apply multilayer networks to characterize patterns of genetic similarity in *var* gene repertoires through time for both a theoretical model and an empirical data set from Ghana, unique in its depth of sampling and coverage over multiple seasons. The patterns of modularity we describe should be relevant to other pathogen systems, whether possessing multi-copy gene families [3] or multiple independent loci encoding different antigens.

## Results

The *var* genes encode the major antigen of the blood stage of infection, PfEMP1 (*Plasmodium falciparum* erythrocyte membrane protein 1) [18]. Besides immune evasion, PfEMP1 promotes adherence of erythrocytes to blood vessels, leading to clinical disease manifestations. Hence *var* genes play an important role in malaria epidemiology. Each parasite genome has a ‘repertoire’ of 50-60 unique *var* gene copies, sequentially expressed to produce different variants of PfEMP1 during infection. In endemic regions of high transmission, *var* genes exhibit enormous diversity [16, 17, 19] resulting from evolutionary innovation at two levels of organization. At the gene level, *var* gene variants can be generated through both mutation and ectopic recombination [20, 21], with tens of thousands of variants documented in local populations [16]. At the repertoire level, variation in gene composition is obtained through sexual (meiotic) recombination in the mosquito vector [20, 22, 23].

Transmission and *var* diversity are positively correlated in the evolutionary history of malaria; that is, areas of low and high transmission have evolved respectively towards low (e.g., South America) and high (e.g., sub-Saharan Africa) diversity. As our questions target systems of high diversity and transmission, and because the results of the two neutral models are qualitatively similar, we focus in the main text on comparing immune selection to complete neutrality for a regime of high diversity, and present results for generalized immunity and for low and medium diversity in the Supporting Results.

We sampled simulated populations monthly for 25 years, with a total of 300 time points, where each time point is a snapshot of changes accumulated in the parasite population during 30 days. We used a temporal multilayer network to analyze population structure in these time series. Competition between repertoires, as well as their evolution and persistence, are intimately interlinked processes emanating from genetic differences. Hence, each layer (time point) was constructed to represent a network of genetic similarity, in which nodes are repertoires and intralayer edges encode genetic similarity between repertoire pairs in terms of their *var* gene composition (Methods; [15]). Using the same metric, we connected layers to each other via unidirectional interlayer edges depicting the genetic similarity between a repertoire in time *t* to those in time *t* + 1 (Fig. 1; Methods). This definition of interlayer edges maps the genetic relationship among repertoires—the process determining persistence—into the network, explicitly creating temporal dependency between layers. We characterized network structure by looking for modules using Infomap, an algorithm that explicitly considers the multilayer structure and the temporal ‘flow’ associated with interlayer edges [24–26]. These modules: (i) contain repertoires that are genetically more similar to each other than to other repertoires in the network; and (ii) are defined within and across layers (across time).

**Figure 1.**
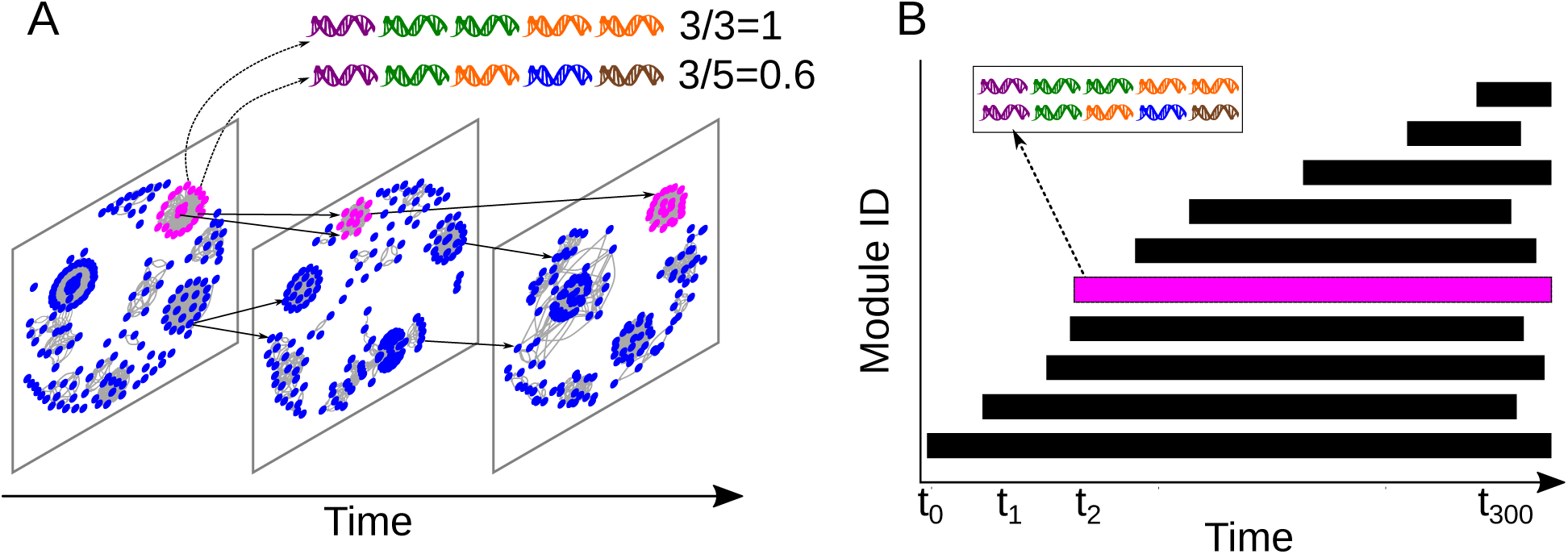
A toy example for a temporal multilayer network of repertoire genetic similarity, and its associated modular structure. In (A), each layer represents a time point. The network in each time point depicts the genetic similarity between pairs of repertoires, where each repertoire is a combination of genes. The measure of genetic similarity is asymmetric to take into account the asymmetric competition resulting from different numbers of distinct gene variants, as shown in the example of the two repertoires: while one repertoire has 3 unique *var* genes out of 5 (depicted as DNA symbols with different colors), the other has 5 unique *var* genes and so will outcompete the first one. Interlayer edges between repertoires leading from any time point *t* to time point *t* + 1 were defined in the same way, but these links only point in one direction to represent temporal flow. For clarity, we present only a few interlayer edges (in black). (B) We used an algorithm for community detection in networks to identify ‘modules’ (depicted as rectangles), which contain repertoires that are more genetically similar to each other than to repertoires from other modules. Modules persist in time but the number and identity of the repertoires which compose them can change (e.g., repertoires can be removed by host immunity and can appear as a result of recombination). New modules appear in time, while others die out. Pink repertoires in (A) are part of the same, pink, module in (B).

## The temporal and non-neutral population structure

Under both immune selection and complete neutrality, we find a dynamic modular structure, in which modules are constantly generated and die out. Interestingly, the structure generated under the former exhibits long-lived coexisting modules, whereas the one generated under the latter is characterized by short-lived ones (Fig. 2A,B). We can examine the role of immune selection by comparing the persistence of repertoires to that of modules. When immune selection is at play, we find that modules persist longer than the repertoires that compose them (Fig. 2C). This mismatch in persistence indicates that modules are not necessarily composed of the same repertoires across time and that repertoires do not necessarily have to persist as long as the module they occupy. By contrast, in the neutral scenarios, modules are short-lived compared to their repertoires (Fig. 2D; Fig. S2). This contrast indicates that the structure of modules in the presence of immune selection is a result of the stabilizing competition between repertoires, which pushes them to be as dissimilar as possible. Under neutrality, genetic changes accumulate due to antigenic drift, until reaching a point at which the repertoire population is sufficiently different to create a new set of modules. Immune selection therefore acts as a form of balancing selection, maintaining long-lived modules.

**Figure 2.**
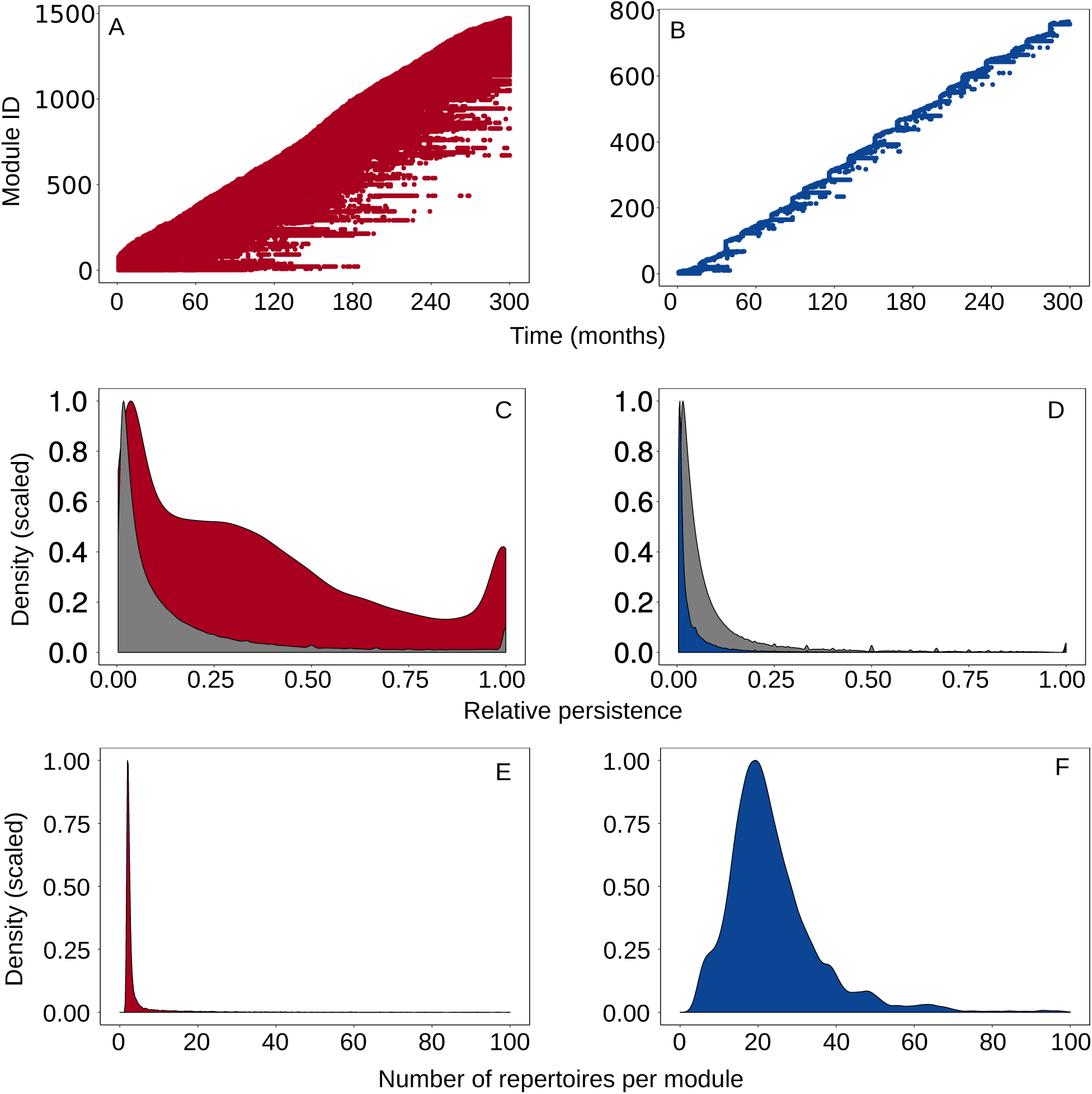
Temporal population structure in the high diversity regime. The left and right columns represent the selection (dark red) and neutral (dark blue) scenarios, respectively. (A,B) Example of population structure from one run of the agent-based model. Each line on the y-axis corresponds to a different module and a circle depicts the occurrence of the module in a given layer. Modules are generated and die out. (C,D) The selection scenario is characterized by modules which persist for much longer than the repertoires that compose them (compare dark red to gray density plots). An opposite trend is evident in the complete neutrality scenario. See Methods for details on how we calculated relative persistence. (E,F) Density plots depicting, *ℛ* the number of repertoires assigned to a module in a given layer.*ℛ* has a theoretical maximum of the number of repertoires in the layer (if there is only one module) and a theoretical minimum of 1 (when each repertoire is assigned to its own module). Immune selection drives repertoires to be as different as possible, resulting in low number of shared genes. Consequently,*ℛ* is low in the immune selection case compared to the neutral scenario. Results in (C-F) are for data pooled across 50 runs of the agent-based model.

Limiting similarity and module coexistence in the selection scenario lead to a high number of modules in any given layer and consequently to a low number of repertoires per module (Fig. 2E). An extremely high proportion of the modules contains only one or two repertoires. By contrast, in the two neutral scenarios, most modules contain about forty repertoires (Fig. 2F; Fig. S2). This is a result of (i) short-lived modules with low tendency to coexist; and (ii) the high number of recombinant repertoires, which, in the absence of immune selection, are not purged and thus fill antigenic space and blur the clear niche separation observed in the selection scenario.

## Comparison to empirical data

We tested our theoretical findings using parasite isolate data collected in an age-stratified longitudinal study in the Bongo District, Ghana [27]. This area is characterized by high seasonality, with a prolonged dry season (Nov-May) and a short wet season (Jun-Oct). Our data set contains four surveys carried out across both the dry and wet seasons; it is the only longitudinal data set available of an asymptomatic *P. falciparum* reservoir population in a given location using deep sequencing.

To validate our theoretical results with empirical data requires generating an expectation benchmark for the population structure under the empirical sampling scheme with the ABM. To this end, we first need to corroborate our theoretical predictions when the seasonality of the sampling area is included. We find that, although seasonality does leave a signature in the formation of modules, the differences between the immune selection and the neutral scenarios remain qualitatively similar (see results and discussion on seasonality in Supporting Results). Next, we ran 100 ABM simulations with parameter values that capture our uncertainty of the exact local parameters in the sampling area, as was done in [15] (Supporting Methods). Our benchmark simulations included, beyond seasonality, the two sessions of indoor residual spraying (IRS) conducted as interventions during the sampling period.

We compared the empirical data to the benchmark simulations across scenarios (Fig. 3A; Fig. S3) by calculating the probability of a module to persist from layer 1 to layer 4 (survival analysis), because we expect that under immune selection modules would persist for longer compared to the neutral scenarios. As expected, module survival probability was higher in the empirical data and under immune selection compared to complete neutrality (Fig. 3B). The similarity between immune selection and generalized immunity is a result of the short time-scale of the empirical data (only 4 layers). Because we match the distribution of infection duration between the generalized immunity and the immune selection scenarios (see Methods), a longer time scale is needed to differentiate them. Hence, on a short time-scale, module persistence under generalized immunity is longer than under complete neutrality but similar to that of immune selection (Fig. S2). Supporting this notion is the finding that networks for empirical data and immune selection differ from those for generalized immunity in structural aspects directly related to persistence, namely the distribution of intra- and inter-layer edge weights and the in- and out-strength of nodes (Fig. S4). This analysis provides evidence for the role of immune selection in maintaining niche coexistence and persistence in an empirical setting.

**Figure 3.**
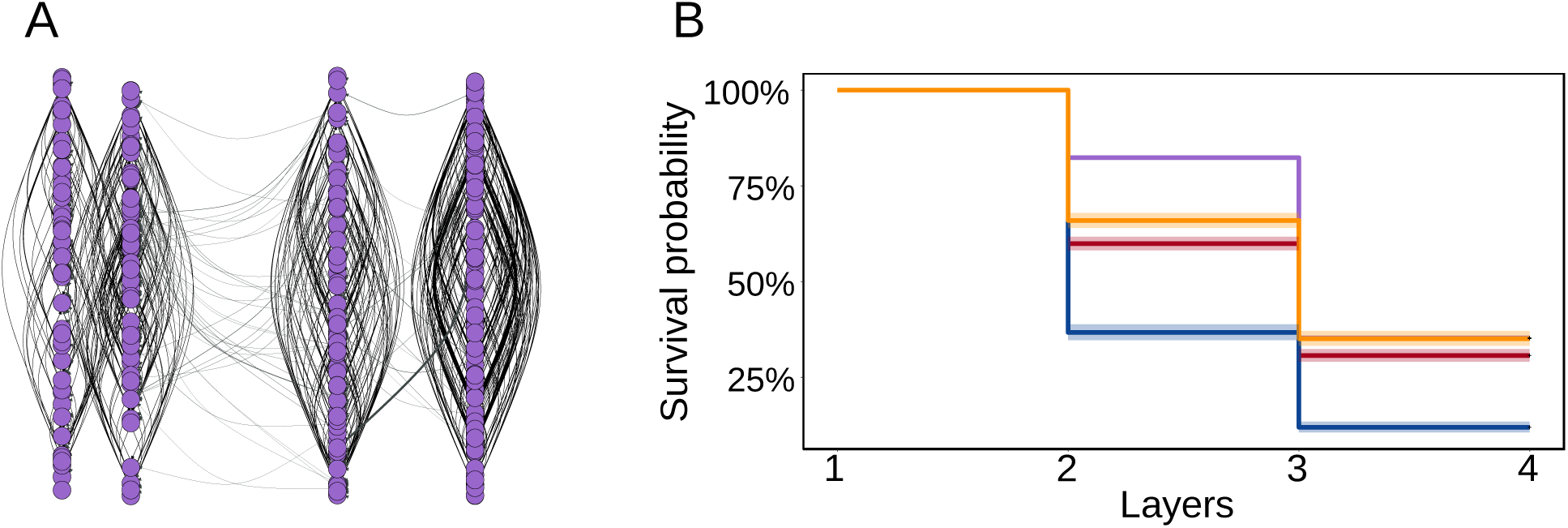
Comparison of empirical data to agent-based model (ABM) simulations. (A) The empirical network (see examples for simulated networks in Fig. S3). The layers are organized by the order of surveys (survey 1 is at the left), and the spacing between them is proportional to the time passed in months between surveys (see Methods and Table S1 for details on the sampling). Within each layer, nodes are ordered randomly. Intra-layer edges are horizontal and depicted in black; interlayer edges are vertical and colored in gray. (B) Analysis of module survival. Curves were calculated with a Kaplan-Meier analysis for the modules that were created in layer 1. The shadowed area represents the 95% confidence intervals across 100 runs of the ABM.

## Epidemiological consequences of strain structure

The modular structure we have uncovered may affect epidemiological parameters because immunity gained to a *var* repertoire from one module facilitates immunity to other repertoires from the same module, and infection with a repertoire from a different module adds exposure largely to *var* genes that have not been seen before. Also, despite modularity, genetic similarity between any two repertoires is generally low because of the very large diversity of the gene pool to begin with. To examine if the modular structure does indeed have an epidemiological meaning, we used the duration of infection as a metric by generating curves that capture the decline in duration of infection with accumulated infections in an individual host. Using the networks produced by the ABM, we simulated infections in naive hosts for one year under the two scenarios, explicitly incorporating the temporal structure and epidemiological force of infection present in the system.

We find that the duration of infection declines at a faster rate when hosts are infected with repertoires originating from the same module, compared to infections with repertoires from different modules (Fig. 4A). Because neutrality also produces modularity, we still observe the difference between the curves but this effect is not as pronounced as under selection (Fig. 4B,C). This is because in the absence of selection, the difference in genetic diversity between repertoires that belong to either the same or different modules is relatively low.

**Figure 4.**
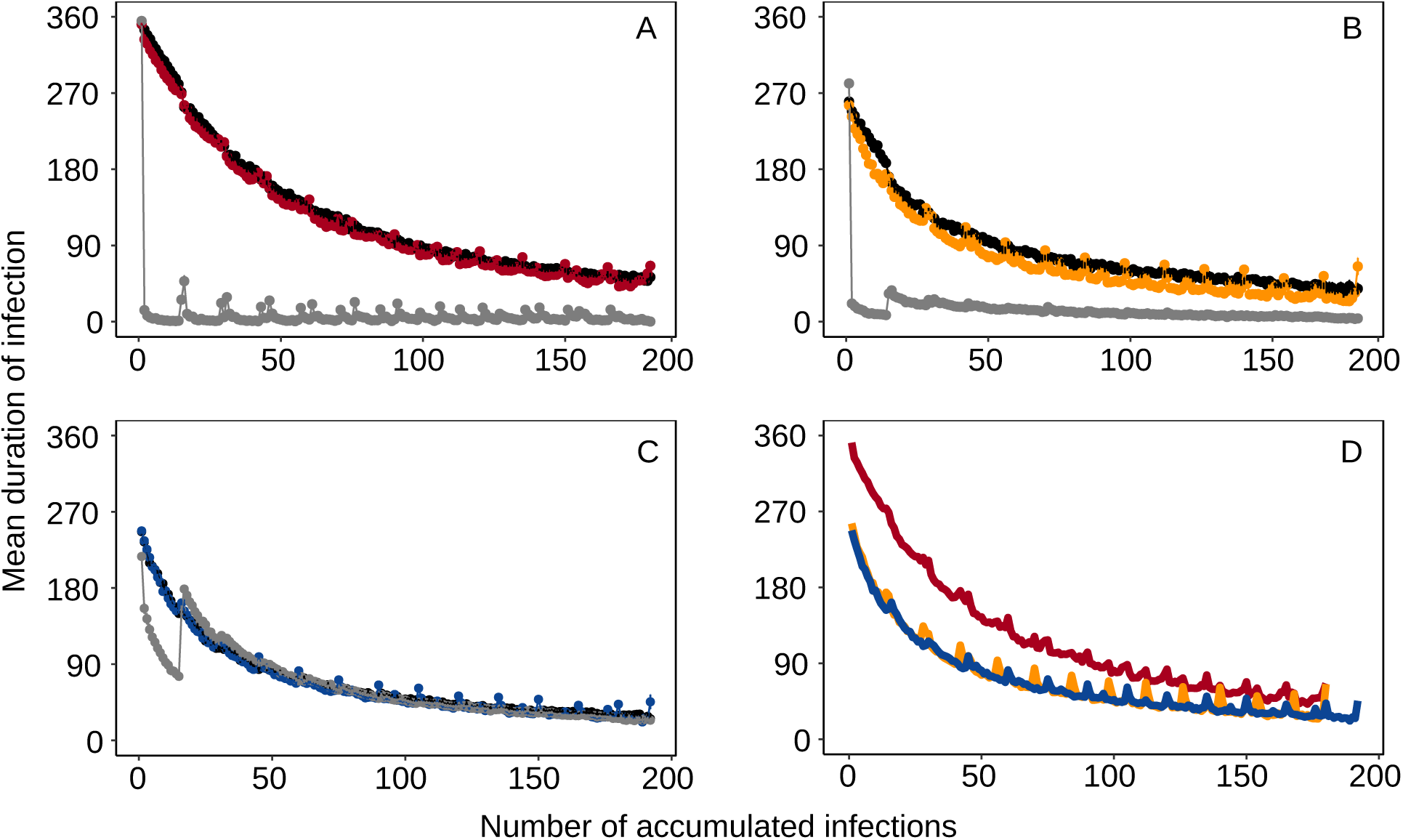
Epidemiological consequences of repertoire population structure under high var gene diversity. The figure shows the decline in duration of infection as a function of the number of infections accumulated in a naive host during one year. Panels (A), (B) and (C) correspond to selection, generalized immunity and complete neutrality, respectively. In any given simulation, a naive host was infected with repertoires originating either from the same module (within-module infections; gray), from different modules (between-module infections; black) or randomly (colored). Each point is the average duration of infection across 20 runs of the agent-based model, for 10 hosts with 5 random starting layers in 10 different modules (see Methods for details on experimental design). We are interested in the comparison of the decreasing trends for the duration of infection across scenarios. The small fluctuations overlaid on these trends reflect intermittent increases in this duration. They are the result of the way infections were sampled over discrete time steps, by advancing from one layer to the next, in the design of the simulations, see Supporting Methods for details). The number of infections in each layer was determined as a function of the entomological inoculation rate of the ABM simulation. (D) A direct comparison of the curves for random infections from panels A-C. The curve of immune selection is above that of the neutral scenarios because repertoires in the neutral scenarios consist of a higher proportion of identical *var* genes, compared to repertoires in the immune selection scenario. In all panels error bars are 95% confidence intervals.

In nature, however, infections can occur with repertoires belonging, or not, to the same module. We therefore additionally sampled repertoires uniformly at random, regardless of their module affiliation. This generates a curve that is very similar to that of the between-module infections because under high diversity most repertoires are classified to their own module within a layer (Fig. 2E,F; Fig. S2). A direct comparison of these curves (Fig. 4D) shows that in populations structured by immune selection, the duration of infection is generally higher than in populations assembled by stochastic processes alone (the red curve is above the other two). This is because (i) immune selection maintains higher *var* gene diversity at the population level; and (ii) competition drives repertoires to be as different from each other as possible, making the genetic diversity within a repertoire higher on average under immune selection.

An exponential fit of the form *d* = *ae*^-*bi*^ (where *d* is the duration of infection and *i* is the number of accumulated infections) to these curves for the first 100 infections indicated about 16% faster decline in *d* in the complete neutrality scenario compared to the immune selection scenario (*b*_*selection*_ = 0.0154 *±* 2.1 · 10^-4^s.e. vs. *b*_*neutral*_ = 0.0183 *±* 4.6 · 10^-4^s.e. and *b*_*generalized*_ = 0.0183 *±* 5.2 · 10^-4^s.e.). These model fits predict a 3-weeks longer infection period under selection compared to neutrality in a 5-months child (i.e., after approximately 75 infections). This underestimation ignores the fact that repertoires in the neutral scenarios consist of a higher proportion of identical *var* genes compared to repertoires in the selection case (expressed as a lower y-intercept for the former (Fig. 4D). When considering this within-repertoire diversity, the model fits predict a 51-day longer infection under selection.

## Discussion

We find that competitive interactions between genomes act as a stabilizing force which dynamically organizes the hyper-diversity of *var* gene repertoires into temporally-persistent modules, akin to emerging niches in the immune space of the human host population. By considering the temporal dimension, we therefore recover the clustered nature characteristic of stabilizing competition/balancing selection, which was not evident in static analyses at high gene diversity [15]. This clustering is also evident on a shorter time scale for field data. Thus, immune selection acts to promote persistence of clusters of gene combinations over epidemiological time scales, in addition to persistence of the genes themselves over evolutionary time [2, 28]. In an ecological context, stabilizing competition between species in ecosystems and OTUs in microbiomes was also shown to promote coexistence and the formation of clusters, albeit for much lower dimensional trait spaces [7–10].

Earlier strain theory described the stable coexistence of genetically-discordant repertoires for lower gene pool sizes [14]. It also postulated that the reproductive number of the parasite *R*_0_ in a given human population is given by the sum of the *R*_0_s of each of the circulating strains [29]. However, it remains unclear what constitutes a ‘strain’ in regions where antigen diversity, transmission and recombination are all high, particularly given that the repertoire population is extremely diverse and dynamic. The continuous genetic coherence, which results in the emergence of unique and persistent niches in antigenic space, suggests that the modules we have identified can play the role of strains. As such, strains are dynamic entities that emerge from the interaction between parasite genomes and the human immune system. Although generated by very different mechanisms, these modules are conceptually similar to viral quasispecies where high mutation rates and negative selection by host immunity create continuously changing repositories of viral variants, which are the source of virus adaptability [30].

The modular structure we find in the immune selection scenario implies a slower acquisition of immunity by individual hosts (compared to neutrality). Duration of infection is a key epidemiological parameter, which together with transmission rate, ultimately determines *R*_0_. Existing epidemiological models of malaria are able to incorporate aspects of repeated infection but neither strain diversity per se nor its overlap structure [31]. How to account for these aspects in transmission models that remain sufficiently parsimonious for epidemiological application remains an open question. Consideration of dynamic strains (or modules) provides one way forward. Incorporating diversity and its structure may prove particularly relevant to modeling interventions. Control efforts can lower diversity either directly by clearing infections (e.g., antimalarial drugs) or through reduction in transmission by targeting the vector (e.g., bed nets or IRS). Our simulations and data clearly show that bottlenecks to transmission (and associated generation of *var* diversity), such as seasonality, limit the persistence of modules, while the replenishment and the restructuring of diversity during the following wet season create new ones (Supporting Results). Immune selection can counter the effects of seasonality and/or interventions by promoting persistence of modules across these bottlenecks. This kind of resilience implies that upon lifting the intervention, the remaining diversity should be able to quickly restructure itself to occupy available niches in immune space. A recent study in Northern Ghana following 7 years of IRS has shown a consistent reduction in entomological inoculation rate (EIR) and sporozoite rates compared to a nearby control site [32]. Yet the same study also showed that upon removing the IRS intervention, EIR increased from 30 to 90 infectious bites per person per month in only two years. Monitoring *var* gene diversity and its structure may provide further insight, as a reduction in epidemiological parameters per se may not be a sufficient condition for elimination. Previous arguments on multiple fixed strains and *R*_0_, coupled with our findings on structure and infection duration, suggest the existence and raise the open question of a threshold in antigenic (and associated genetic) diversity in the transition to elimination.

One possible limitation of the approach taken here is its potential sensitivity to the community detection method we used. While this can, in principle, be true, the patterns we obtain are sensible from a theoretical perspective. Also, on the technical side, there is currently no other method available to perform community detection in directed and weighted networks with directed interlayer edges in large networks. Because quantitative network properties such as the number of modules can be sensitive to the exact parameterization of the algorithm’s implementation, or to the cut-off imposed on network edge weights, we emphasize that we are primarily interested in the qualitative differences between the immune selection and neutral scenarios.

From this qualitative perspective, temporal multilayer networks present an opportunity to extend the scope of current phylodynamic theory, which relates the epidemiology and transmission of pathogens to phylogenetic structure [33, 34]. Because trees are a particular case of networks, the multilayer network approach we devise here can extend traditional phylodynamic analysis to pathogens evolving through recombination. As in phylodynamics, the structure of our network reveals information about the interplay between selective forces and transmission dynamics. There is, in particular, a clear difference in structure between the immune selection and neutral scenarios. We also find differences in network structure between regimes with different diversity, whereby low diversity is characterized by a replacement regime instead of module coexistence (see Supporting Results)—patterns that reflect differences in transmission dynamics and evolutionary rates between these regimes.

Our findings are generally relevant for pathogens characterized by large antigenic spaces. One extreme example is found in *Trypanosoma brucei*, the causal agent of African sleeping sickness. Its genome contains about 1,000 copies of the *vsg* gene, the vast majority of which are pseudogenes. This enormous within-genome diversity provides a clear advantage to the parasite, allowing it to ‘assemble’ new active genes from pseudogenes via gene conversion, once it has exhausted its functional copies during an infection [35]. Other examples are found in the fungus *Pneumocystis carinii* and in the bacterium *Neisseria gonorrhoeae* [3]. Beyond multicopy family genes, the theory and analyses we have developed should also apply to sets of genes encoding different antigens.

Competition for hosts is a powerful feature of parasites’ life history, enabling the emergence and maintenance of a dynamical strain structure across bottlenecks of transmission, and affecting the rate at which host immunity is gained. For malaria, understanding the consequences of parasite population structure for elimination constitutes an important direction for future research.

## Methods

### Agent-based model

The ABM is flexible enough to include seasonality and methods for malaria control to compare simulated dynamics to empirical data (see below). We used the same ABM implementation as developed by [15]. A complete description of the model can be found there. Here, we briefly describe the main elements of the model.

#### Structure of human and parasite populations

The human population in the ABM has a fixed size of *H* = 10, 000, with a given age structure. Hence, whenever a host dies, another is born. The genome of one individual *P. falciparum* parasite was considered as a repertoire consisting of a set of 60, not necessarily unique, *var* genes, *ℛ* =*{g*_1_, *g*_2_, …, gl*}* where 1*≤l≤*60 is the index of the gene within the repertoire and *g∈U* [1, *G*] is the ID of the gene in a pool of *G* genes. Each *var* gene was composed of two variants (hereafter, alleles), corresponding to two epitopes. This model of *var* genes is supported by empirical evidence for two hypervariable regions bounded by three semi-conserved regions [36, 37]. Hence, each gene is defined as 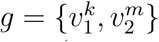where the superscripts *k, m ∈ U* [1, *A*] are the ID of the allele in a pool of *A* alleles (the two epitopes have separate pools which are equal in size), and the subscript defines the epitope. A focal population of the parasite is initialized from an external gene pool representing regional diversity, from which immigration also occurs (at a rate of one repertoire per day). Each simulation starts with 20 infected hosts, each with one repertoire selected at random from the pool.

#### Transmission dynamics

Vectors (mosquitoes) are not explicitly modeled. Instead, we set a biting rate *b* so that the average waiting time to the next biting event is equal to 1*/*(*b×H*). When a biting event occurs, two hosts are randomly selected, one donor and one recipient. If the donor has infectious parasite repertoires, the receiver will be infected with a probability of *p* (i.e., transmission probability). If the donor is infected with multiple repertoires in the blood stage, then the transmission probability of each strain is *p/I*, where *I* is the number of repertoires.

#### Mechanisms of genetic change

During the sexual stage of the parasite (within mosquitoes), different parasites can exchange *var* repertoires through meiotic recombination. The receiver host can receive either recombinant repertoires, original repertoires or a combination. During the asexual stage of the parasite (blood stage of infection), *var* genes within the same repertoire can exchange epitope alleles through mitotic (ectopic) recombination (at a rate of *ρ* = 1.87*×*10^-7^ per day). Also, epitopes can mutate (at a rate of *µ* = 1.42 *×*10^-8^ per day). Mitotic recombination, mutation and immigration generate new *var* genes [21].

## Within-host dynamics

Each repertoire is individually tracked through its entire life cycle, encompassing the liver stage, asexual blood stage, and the transmission and sexual stages. Because we do not explicitly model mosquitoes, we delay the expression of each strain in the receiver host by 14 days to account for the time required for the sexual stage (in the mosquito) and the liver stage (in the host). Specifically, the infection of the host is delayed 7 days to account for the time required for gametocytes to develop into sporozoites in mosquitoes. When a host is infected, the parasite remains in the liver stage for additional days before being released as merozoites into the bloodstream, invading red blood cells and starting the expression of the *var* repertoire.

During the expression of the repertoire, the host is considered infectious with the active repertoire. The expression order of the repertoires is randomized for each infection, while the deactivation rates of the *var* genes is controlled by the host immunity. When one gene is actively expressed, host immunity ‘checks’ whether it has seen any of its epitopes in the infection history. In the immune selection model, the deactivation rate changes so that the duration of active period of a gene is proportional to the number of unseen epitopes. After the gene is deactivated, the host adds its deactivated alleles to the immunity memory. A new gene from the repertoire then becomes immediately active. The repertoire is cleared from the host when the whole repertoire of *var* genes is depleted. The immunity towards a certain epitope wanes at a rate *w* = 1*/*1000 per day [38].

## Scenarios

In the scenario of immune selection, the duration of infection in a host depends on the history of infection with given *var* genes (or their alleles). The model of ‘complete neutrality’ retains the process of transmission but does not consider any aspect of infection history. Hence, individuals clear infection after a given amount of time (matched to the average duration of infection from the selection scenario). In the neutral model of ‘generalized immunity’, the duration of infection depends on the number of infections, but not on their specific genetic composition. To parameterize this model, we matched the relationship between the duration of infection and accumulated number of infections from the selection scenario [15]. In both neutral models we ran the exact same Agent-Based Model (ABM) implementation of the immune selection scenario, conserving all its parameters except for the competitive interactions and the resulting duration of infection. Because we know that in nature hosts build immunity to malaria infections, the neutral scenarios can be regarded as process-based null models against which we test the structure obtained from simulations with immune selection and the empirical data. For each combination of diversity regime (low, medium, high) and immune scenario (immune selection, complete neutrality and generalized immunity), we analyzed data generated by 50 simulations of the ABM.

## Construction of temporal networks

We calculated genetic similarity of repertoire *i* to repertoire *j* as *S*_*ij*_ = (*N*_*i*_ *∩ N*_*j*_)*/N*_*i*_, where *N*_*i*_ and *N*_*j*_ are the number of unique alleles for repertoires *i* and *j*, respectively (the genetic similarity of repertoire *j* to repertoire *i* was calculated as *S*_*ji*_ = (*N*_*i*_*∩N*_*j*_)*/N*_*j*_). We used a directional metric because of the asymmetric competition resulting from different numbers of unique alleles in a repertoire [15]. We used the same metric (*S*) for both intra- and inter-layers because it situates the inter- and intra-layer edges on the same scale, which is crucial when looking for optimal partitioning in multilayer networks [39,40]. To optimize the signal-to-noise ratio in our analysis, we imposed a cut-off on edge weights (see Supporting Methods).

## Community detection

To capture the organization of the population into groups of highly similar repertoires we used the map equation as an objective function to calculate the optimal partition of the network. Briefly, the map equation is a flow-based and information-theoretic method (implemented with Infomap), recently extended to multilayer networks, which calculates network partitioning based on the movement of a random walker on the network (see [24–26] for details). In any given partition of the network, the random walker moves across nodes with proportion to the direction and weight of the edges. Hence, it will tend to stay longer in dense areas where there are many repertoires similar to each other. These areas can be defined as ‘modules’. The time spent in each module can be converted to an information-theoretic currency using an objective function called the map equation. The best network partition corresponds to that with the minimum value of the map equation [24, 25]. This method has been applied to describe temporal flow in networks that do not have interlayer edges [41]. In our particular network, once the random walker moves along an interlayer edge, it cannot go back, capturing the temporal flow in the network.

## Quantification of structure

We calculated *ℛ*, the average number of repertoires per module in each layer. The theoretical maximum of *ℛ* is the number of repertoires in the layer (if there is only one module) and its theoretical minimum is 1 (when each repertoire is assigned to its own module). Values close to the minimum indicate that repertoires are at the ‘limit’ of their limiting similarity, as each repertoire is as different from any another as possible. We calculated the relative persistence of repertoires and modules as *𝒫* = (*l*_*d*_ *- l*_*b*_)*/*(*l*_*max*_ *- l*_*b*_), where *l*_*b*_ and *l*_*d*_ refer to the layers in which a module (or repertoire) first appeared or died, respectively and *l*_*max*_ is the maximum number of layers (*l*_*max*_ = 300 in our simulations). This metric has a maximum theoretical value of 1 when a module (repertoire) persists from the layer when it was born till *l*_*max*_ and a theoretical minimum of 1*/*(*l*_*max*_*-l*_*b*_) when a module (repertoire) persists for a single layer. We also examined the results for absolute persistence (i.e., the number of layers), and these do not qualitatively change the conclusions.

## Empirical data

The empirical data used was collected from a study performed across two catchment areas in Bongo District (BD), Ghana located in the Upper East Region near the Burkina Faso border. This age-stratified serial cross-sectional study was conducted across four surveys: the end of the wet season, October 2012 (S1), the end of the dry season between May-June 2013 (S2), the end of the wet season, October 2013 (S3), and the end of the dry season May-June 2014 (S4). Each survey lasted approximately 3 to 4 weeks. During the sampling period, two rounds of IRS intervention were performed (between October-December 2013 and between May-July 2014). We sequenced and genotyped the *var* DBL*α* domain, which is a reliable marker for differentiating *var* genes. DBL*α* reads were clustered at 96% pairwise identity and translated into all six reading frames and classified into either upsA or upsB/upsC (i.e., non-upsA) groups [19]. Details on the study population, data collection procedures, molecular and genetic work, and epidemiology have been published elsewhere [15,27]. While data from S1 and S2 were used in these two previous studies, data from S3 and S4 are used here for the first time. A summary of the data across surveys can be found in Table S1.

The study was reviewed and approved by the ethics committees at the Navrongo Health Research Centre, Ghana; Noguchi Memorial Institute for Medical Research, Ghana; University of Melbourne, Australia; and the University of Chicago, United States.

## Comparison of empirical and simulated benchmark networks

Following [15], we constructed each layer in the empirical network based on pairwise similarities of unique upsB/upsC DBL*α* types because upsA genes tend to be more conserved and thus have disproportionately higher sharing rates among repertoires [19]. Considering all genes does not qualitatively change the results. Because sequencing did not provide information on distinct variants in the DBL*α* epitopes, we considered whole DBL*α* instead of variants as the unit of similarity [15]. Since infections with multiple parasite genomes (multiplicity of infection; MOI*>* 1) are very common in malaria endemic regions, we selected isolates with a total number of upsB/upsC DBL*α* types ranging from 40-55 copies, to maximize the probability of selecting hosts with single-genome infections (MOI=1). This resulted in 97, 67, 66 and 50 isolates in layers 1-4, respectively. Edge weights were defined using the same metric and cutoff (90%) as in the theoretical work. We compared this network to simulated networks of equivalent size. To account for the uneven sampling periods between the four surveys, we rescaled the interlayer edge weights in the empirical and simulated networks. Details in the Supporting Methods.

## Simulations of host infection

We simulated repeated infections of naive hosts with repertoires originating in the same module, in different modules or regardless of module. We compared between curves that depicted the decrease in the duration of infection as a function of the number of accumulated infections in each of these dynamics. The infection takes place within a layer for a given number of bites, which depend on the force of infection, and then moves on to the next layer. This explicitly considers the temporal component. Details on the simulations are in the Supporting Methods.

## Data and code

THe following code is available at GitHub: The original C++code for the ABM. R code for analysis. Python code for the sequence cleaning pipeline. Python code to determine DBL*α* types. Sequences have been deposited at DDBJ/ENA/GenBank under the Bio-Project Number: PRJNA 396962.

## Acknowledgments

This research was supported by the Fogarty International Center at the National Institutes of Health [Program on the Ecology and Evolution of Infectious Diseases (EEID), Grant number: R01-TW009670]. S.P. was supported by a James S. McDonnell Foundation 21^st^ Century Science Initiative – Postdoctoral Program in Complexity Science-Complex Systems Fellowship Award and by a Fulbright Fellowship from the U.S. Department of State. We thank Martin Rosvall and Daniel Edler for advice and support with Infomap. We wish to thank the participants, communities and the Ghana Health Service in Bongo District Ghana for their willingness to participate in this study. We would also like to thank the personnel at the Navrongo Health Research Centre for sample collection and parasitological assessment/expertise. We are grateful to Abraham R. Oduro, Anita Ghansah, and Kwadwo Koram for their helpful input related to the field study, to Gerry Tonkin-Hill for the development of the Illumina sequencing cleaning and clustering pipelines, and to Michael F. Duffy for insightful comments on an earlier version of the manuscript. We appreciate the support of the University of Chicago through computational resources at the Midway cluster.

## References

[1] Charlesworth D. Balancing selection and its effects on sequences in nearby genome regions. Plos Genet. 2006;2(4):e64.

[2] Larremore DB, Sundararaman SA, Liu W, Proto WR, Clauset A, Loy DE, et al. Ape parasite origins of human malaria virulence genes. Nat Commun. 2015;6:8368. doi:10.1038/ncomms9368.

[3] Deitsch KW, Lukehart SA, Stringer JR. Common strategies for antigenic variation by bacterial, fungal and protozoan pathogens. Nat Rev Microbiol. 2009;7(7):493–503. doi:10.1038/nrmicro2145.

[4] Figueroa F, Günther E, Klein J. MHC polymorphism pre-dating speciation. Nature. 1988;335(6187):265–267. doi:10.1038/335265a0.

[5] Chesson P. Mechanisms of maintenance of species diversity. Annu Rev Ecol Syst. 2000;31(1):343–366.

[6] Adler PB, Hillerislambers J, Levine JM. A niche for neutrality. Ecol Lett. 2007;10(2):95–104. doi:10.1111/j.1461-0248.2006.00996.x.

[7] Scheffer M, van Nes EH. Self-organized similarity, the evolutionary emergence of groups of similar species. Proc Natl Acad Sci U S A. 2006;103(16):6230–6235. doi:10.1073/pnas.0508024103.

[8] Jeraldo P, Sipos M, Chia N, Brulc JM, Dhillon AS, Konkel ME, et al. Quantification of the relative roles of niche and neutral processes in structuring gastrointestinal microbiomes. Proc Natl Acad Sci U S A. 2012;109(25):9692–9698. doi:10.1073/pnas.1206721109.

[9] D’Andrea R, Ostling A. Can Clustering in Genotype Space Reveal “Niches”? Am Nat. 2016;187(1):130–135. doi:10.1086/684116.

[10] Rocha RD, Riolo M, Ostling AM. Competition and immigration lead to clusters of similar species, not trait separation; 2018.

[11] Gupta S, Maiden MCJ, Feavers IM, Nee S, May RM, Anderson RM. The maintenance of strain structure in populations of recombining infectious agents. Nat Med. 1996;2(4):437–442. doi:10.1038/nm0496-437.

[12] Gupta S, Day KP. A theoretical framework for the immunoepidemiology of Plasmodium falciparum malaria. Parasite Immunol. 1994;16(7):361–370.

[13] Gupta S, Anderson RM. Population structure of pathogens: The role of immune selection. Parasitol Today. 1999;4758(1994):497–501. doi:10.1016/S0169-4758(99)01559-8.

[14] Artzy-Randrup Y, Rorick MM, Day KP, Chen D, Dobson AP, Pascual M. Population structuring of multi-copy, antigen-encoding genes in Plasmodium falciparum. Elife. 2012;2012(1):e00093. doi:10.7554/eLife.00093.

[15] He Q, Pilosof S, Tiedje KE, Ruybal-Pesántez S, Artzy-Randrup Y, Baskerville EB, et al. Networks of genetic similarity reveal non-neutral processes shape strain structure in Plasmodium falciparum. Nat Commun. 2018;9(1):1817. doi:10.1038/s41467-018-04219-3.

[16] Day KP, Artzy-Randrup Y, Tiedje KE, Rougeron V, Chen DS, Rask TS, et al. Evidence of strain structure in Plasmodium falciparum var gene repertoires in children from Gabon, West Africa. Proceedings of the National Academy of Sciences. 2017; p. 201613018. doi:10.1073/pnas.1613018114.

[17] Rorick MM, Artzy-Randrup Y, Ruybal-Pesántez S, Tiedje KE, Rask TS, Oduro A, et al. Signatures of competition and strain structure within the major blood-stage antigen of Plasmodium falciparum in a local community in Ghana. Ecol Evol. 2018;doi:10.1002/ece3.3803.

[18] Kirkman LA, Deitsch KW. Antigenic variation and the generation of diversity in malaria parasites. Curr Opin Microbiol. 2012;15(4):456–462. doi:10.1016/j.mib.2012.03.003.

[19] Ruybal-Pesántez S, Tiedje KE, Tonkin-Hill G, Rask TS, Kamya MR, Greenhouse B, et al. Population genomics of virulence genes of Plasmodium falciparum in clinical isolates from Uganda. Sci Rep. 2017;7(1):11810. doi:10.1038/s41598-017-11814-9.

[20] Freitas-Junior LH, Bottius E, Pirrit LA, Deitsch KW, Scheidig C, Guinet F, et al. Frequent ectopic recombination of virulence factor genes in telomeric chromosome clusters of P. falciparum. Nature. 2000;407(6807):1018–1022. doi:10.1038/35039531.

[21] Claessens A, Hamilton WL, Kekre M, Otto TD, Faizullabhoy A, Rayner JC, et al. Generation of Antigenic Diversity in Plasmodium falciparum by Structured Re-arrangement of Var Genes During Mitosis. Plos Genet. 2014;10(12):e1004812. doi:10.1371/journal.pgen.1004812.

[22] Paul RE, Packer MJ, Walmsley M, Lagog M, Ranford-Cartwright LC, Paru R, et al. Mating patterns in malaria parasite populations of Papua New Guinea [see comments]. Science. 1995;269(September):1709–1711.

[23] Su X, Ferdig MT, Huang Y, Huynh CQ, Liu A, You J, et al. A genetic map and recombination parameters of the human malaria parasite Plasmodium falciparum. Science. 1999;286(5443):1351–1353.

[24] Rosvall M, Bergstrom CT. Maps of random walks on complex networks reveal community structure. Proc Natl Acad Sci U S A. 2008;105(4):1118–1123. doi:10.1073/pnas.0706851105.

[25] Rosvall M, Axelsson D, Bergstrom CT. The map equation. Eur Phys J Spec Top. 2010;178(1):13–23. doi:10.1140/epjst/e2010-01179-1.

[26] De Domenico M, Lancichinetti A, Arenas A, Rosvall M. Identifying modular flows on multilayer networks reveals highly overlapping organization in interconnected systems. Phys Rev X. 2015;5(1):011027. doi:10.1103/PhysRevX.5.011027.

[27] Tiedje KE, Oduro A, Agongo G, Anyorigiya T, Azongo D, Awine T, et al. Seasonal Variation in the Epidemiology of Asymptomatic Plasmodium falciparum Infections Across Two Catchment Areas in Bongo District, Ghana. Am J Trop Med Hyg. 2017;97(1):199–212. doi:10.4269/ajtmh.16-0959.

[28] Zilversmit MM, Chase EK, Chen DS, Awadalla P, Day KP, McVean G. Hypervariable antigen genes in malaria have ancient roots. BMC Evol Biol. 2013;13(1):110. doi:10.1186/1471-2148-13-110.

[29] Gupta S, Day KP. A strain theory of malaria transmission. Parasitol Today. 1994;446(I 993):3737–3742. doi:10.1016/0169-4758(94)90160-0.

[30] Domingo E, Sheldon J, Perales C. Viral quasispecies evolution. Microbiol Mol Biol Rev. 2012;76(2):159–216. doi:10.1128/MMBR.05023-11.

[31] Mandal S, Sarkar RR, Sinha S. Mathematical models of malaria–a review. Malar J. 2011;10:202. doi:10.1186/1475-2875-10-202.

[32] Coleman S, Dadzie SK, Seyoum A, Yihdego Y, Mumba P, Dengela D, et al. A reduction in malaria transmission intensity in Northern Ghana after 7 years of indoor residual spraying. Malar J. 2017;16(1):324. doi:10.1186/s12936-017-1971-0.

[33] Grenfell BT, Pybus OG, Gog JR, Wood JLN, Daly JM, Mumford JA, et al. Unifying the Epidemiological and Evolutionary Dynamics of Pathogens. Science. 2004;303.

[34] Volz EM, Koelle K, Bedford T. Viral phylodynamics. Plos Comput Biol. 2013;9(3):e1002947. doi:10.1371/journal.pcbi.1002947.

[35] Taylor JE, Rudenko G. Switching trypanosome coats: what’s in the wardrobe? Trends Genet. 2006;22(11):614–620. doi:10.1016/j.tig.2006.08.003.

[36] Bull PC, Buckee CO, Kyes S, Kortok MM, Thathy V, Guyah B, et al. Plasmodium falciparum antigenic variation. Mapping mosaic var gene sequences onto a network of shared, highly polymorphic sequence blocks. Mol Microbiol. 2008;68(6):1519–1534. doi:10.1111/j.1365-2958.2008.06248.x.

[37] Rask TS, Hansen Da, Theander TG, Pedersen AG, Lavstsen T. Plasmodium falciparum Erythrocyte Membrane Protein 1 Diversity in Seven Genomes - Divide and Conquer. Plos Comput Biol. 2010;6(9):e1000933. doi:10.1371/journal.pcbi.1000933.

[38] Collins WE, Warren M, Skinner JC, Fredericks HJ. Studies on the relationship between fluorescent antibody response and ecology of malaria in Malaysia. Bull World Health Organ. 1968;39(3):451–463.

[39] Mucha PJ, Richardson T, Macon K, Porter MA, Onnela JP. Community structure in time-dependent, multiscale, and multiplex networks. Science. 2010;328(5980):876–878. doi:10.1126/science.1184819.

[40] Pilosof S, Porter MA, Pascual M, Kéfi S. The multilayer nature of ecological networks. Nature Ecology & Evolution. 2017;1:0101. doi:10.1038/s41559-017-0101.

[41] Rosvall M, Bergstrom CT. Mapping change in large networks. Plos One. 2010;5(1):e8694. doi:10.1371/journal.pone.0008694.

